# Feeding a low-carbohydrate and high-protein diet diminishes working memory in healthy mice: possible involvement of miR-539-3p/*Lrp6*/*Igf1r* axis in the hippocampus

**DOI:** 10.1101/2024.04.09.588802

**Authors:** Takeru Shima, Hayate Onishi, Chiho Terashima

## Abstract

A low-carbohydrate and high-protein (LC-HP) diet demonstrates favorable impacts on metabolic parameters, albeit it leads to a decline in hippocampal function among healthy mice. The reduction in working memory induced by LC-HP diets is attributed to the decreased expression of hippocampal IGF-1 receptor (IGF-1R). However, the precise mechanisms underlying this phenomenon remain unexplored. Here, we investigated that by analyzing alterations in hippocampal miRNA profiles. C57BL/6 mice were divided into the LC-HP diet-fed group (25.1% carbohydrate, 57.2% protein, and 17.7% fat as percentages of calories) and the control diet-fed group (58.9% carbohydrate, 24.0% protein, and 17.1% fat as percentages of calories). After four weeks, all mice underwent the Y-maze test, followed by analyses of mRNA and miRNA expressions in the hippocampus. Our investigation revealed that feeding the LC-HP diet suppressed working memory function and hippocampal *Igf1r* mRNA levels in mice. Sequencing of miRNA demonstrated 17 upregulated and 27 downregulated miRNAs in the hippocampus of LC-HP diet-fed mice. Notably, upregulation of miR-539-3p, predicted to modulate *Igf1r* gene expression, was observed. Consequently, we found decreased hippocampal mRNA levels of low-density lipoprotein receptor-related protein 6 (*Lrp6*), a gene modulated by miR-539-3p, in mice fed the LC-HP diet. Furthermore, a significant positive correlation was observed between *Lrp6* and *Igf1r* mRNA levels in the hippocampus. These findings suggest that LC-HP diets may suppress hippocampal function via the miR-539-3p/*Lrp6*/*Igf1r* axis, offering a potential target for nutritional strategies to preserve hippocampal health.

## Introduction

The nutritional composition of diets plays a pivotal role in dietary approaches aimed at preserving both physical and mental health in humans. While prior research has highlighted the advantages of low-carbohydrate and high-protein (LC-HP) diets in enhancing glycemic control [1,2], there are also documented adverse effects, including atherosclerosis and dysfunction in kidneys and vessels [3–5]. Of particular interest are the effects of such diets on brain function: while LC-HP diets have shown benefits in obese individuals [6], they have been associated with diminished hippocampal memory function in healthy animals [7]. This observation holds significance, given the importance of maintaining hippocampal function for overall human well-being. Consequently, unraveling the underlying mechanisms behind the LC-HP diet-induced decline in hippocampal function is imperative and holds considerable relevance in devising appropriate dietary strategies.

A recent investigation proposes a possible involvement of IGF-1 receptor (IGF-1R) with LC-HP diets-induced hippocampal dysfunction [7]. IGF-1R plays a pivotal role in regulating hippocampal neurogenesis [8,9], a critical morphological adaptation essential for maintaining hippocampal function [10,11]. Notably, deletion of *Igf1r* gene has been shown to downregulate hippocampal neurogenesis [12]. A previous study has shown that feeding an LC-HP diet leads to decreased hippocampal mRNA levels of *Igf1r* and *Dcx* [7], a marker of newborn immature neurons [13,14]. Conversely, there is no observed difference in hippocampal mRNA levels of *Bdnf*, the other positive regulator of hippocampal neurogenesis [15,16], between the mice fed an LC-HP diet and those fed a control diet [7]. Thus, it is suggested that LC-HP diet-induced hippocampal dysfunction may stem from dysregulation of IGF-1R-related neurogenesis in the hippocampus. However, the precise mechanisms underlying the downregulation of hippocampal expressions in IGF-1R during LC-HP diet feeding remain to be elucidated.

The current study focused on alterations in microRNA (miRNA) expressions, which are short non-coding RNAs typically ranging from 21 to 25 nucleotides in length [17]. MiRNAs are known to function by binding to complementary segments of messenger RNAs (mRNAs), subsequently leading to the degradation or inhibition of mRNA translation [18]. Their roles in brain development and functions have been documented [19–21], and changes in miRNA profiles are recognized as important regulators and biomarkers for dietary effects [22–24]. Moreover, variations in the nutritional composition of diets have been shown to influence miRNA profiles in various organs and circulation [25–28]. Previous studies have indicated that IGF-1R expression can be regulated by specific miRNAs, including miR-96, miR-181b, and miR-223 [29–31]. Specifically, an upregulation of miR-96 has been associated with decreased hippocampal IGF-1R levels and consequent memory dysfunction [29]. While there is a possibility that these miRNAs may be implicated in LC-HP diet-induced hippocampal dysfunction and the subsequent decrease in hippocampal *Igf1r* mRNA levels, the specific changes in hippocampal miRNA profiles resulting from LC-HP diet consumption remain unclear.

Here, we tested the effects of feeding LC-HP diets over a span of four weeks on hippocampal working memory in healthy mice and the alterations in hippocampal miRNA profiles using miR-Seq to gain insight into the modulation of hippocampal IGF-1R expression.

## 2. Materials and Methods

### 2.1. Animals

Eight-week-old male C57BL/6 mice obtained from SLC Inc. (Shizuoka, Japan) were housed in a temperature-controlled facility set at 21-23□ and maintained on a 12-hour light/dark cycle (lights on from 7:00 to 19:00). During the acclimatization period, the mice had access to a standard pellet diet (Rodent Diet CE-2, CLEA Japan Inc., Tokyo, Japan) and water ad libitum. The experimental procedures were pre-approved (approval No. 23-050) and conducted in accordance with the guidelines established by the Gunma University Animal Care and Experimentation Committee.

### 2.2. Experimental design

Following one week of acclimatization, the mice were divided into two groups based on matching body weights: a group receiving the LC-HP diet (25.1% carbohydrate, 57.2% protein, and 17.7% fat as percentages of calories; n = 10) and a group receiving the control diet (58.9% carbohydrate, 24.0% protein, and 17.1% fat as percentages of calories; n = 10). The diets were obtained from CLEA Japan Inc. (Tokyo, Japan), and their compositions are delineated in Table 1. All mice had access to water ad libitum throughout the four-week experimental period. Subsequently, after the four-week feeding regimen, all mice underwent the Y-maze test.

**Table 1.** Nutritional composition of the diets.

| Ingredients | % |  |
| --- | --- | --- |
|  | Control | LC-HP |
| Cornstarch<br>(including alpha-cornstarch) | 46.2 | 11.9 |
| Casein | 24.5 | 58.8 |
| Sucrose | 10.0 | 10.0 |
| Mineral mix | 7.0 | 7.0 |
| Corn oil | 6.0 | 6.0 |
| Cellulose | 3.0 | 3.0 |
| Cellulose powder | 2.0 | 2.0 |
| Vitamin mix | 1.0 | 1.0 |
| Choline chloride | 0.3 | 0.3 |

LC-HP: Low-carbohydrate and high-protein diet.

### 2.3. Evaluation of working memory

The Y-maze test was conducted following previously described [7]. Prior to the test, mice underwent a three-day acclimation period in a soundproof room for 30 minutes daily. Subsequently, each mouse was placed in the maze’s center and allowed to freely explore the maze (with each arm measuring 43 cm in length, 4 cm in width, and 12 cm in wall height, O’hara & Co., Ltd., Japan). The sequence and frequency of entries into the arms were recorded over eight minutes using a video tracking system (O’hara & Co., Ltd., Japan). The correct ratio for each mouse was calculated when it consecutively navigated through all three arms without revisiting any previously entered arms (the correct ratio = the number of correct trials/[the total number of arm entries - 2]).

### 2.4. Tissue preparation

Two days after the Y-maze test, mice were anesthetized with isoflurane (30% isoflurane in propylene glycol; Dainippon Sumitomo Pharma Co., Osaka, Japan), and the blood samples were obtained from mice by cardiac puncture. And then, the hippocampus was collected in RNAlater™ Stabilization Solution (Invitrogen™). The hippocampus samples were stored at -20□ for subsequent biochemical analysis.

### 2.5. Blood glucose and **β**-ketone assay

Blood glucose and β-ketone levels were measured by the FreeStyle Precision Neo meter (FreeStyle Libre, Abbot, Japan).

### 2.6. Real-time PCR

Total RNA was extracted from the hippocampus tissue using RNeasy Mini Kit with DNase I treatment (Qiagen Inc., USA) according to the protocol provided. RNA quantification was carried out using the Qubit 4.0 (Invitrogen™, USA), and then 1000 ng of RNA was reverse transcribed to cDNA with GeneAce cDNA Synthesis Kit (NIPPON Genetics Co., Ltd., Japan). After that, we measured the mRNA levels of target genes using 5.0 ng of cDNA, primers for each target gene, and PowerTrack™□ SYBR™□ Green Master Mix in StepOne Plus Real-Time PCR 96-well system (Thermo Fisher Scientific Inc., USA). The sequences of primers (forward and reverse) are shown in Supplementary Table 1. The relative levels of each mRNA were calculated by the ΔΔCT method and normalized by β-actin mRNA levels.

### 2.7. miRNA isolation and NGS of miRNA

We used the miRNeasy Micro Kit (Qiagen, Inc., Valencia, CA, USA) following the manufacturer’s instructions for miRNA isolation from the hippocampus. The libraries for miRNA sequencing were prepared utilizing 100.0 ng of RNA, QIAseq miRNA Library Kit, and QIAseq miRNA NGS 48 Index IL (Qiagen Inc., Valencia, CA, USA), according to the manufacturer’s protocol. The library quality was evaluated by measuring the library size in base pairs using the Agilent High Sensitivity DNA Kit and Agilent Bioanalyzer (Agilent Technologies, Santa Clara, CA). During the library construction, each miRNA molecule was tagged with a unique molecular identifier (UMI). Single-end sequencing of 86-bp reads was performed on the NextSeq 500 using the NextSeq 500/550 High Output Kit v2.5 (Illumina Inc., San Diego, CA, USA). The sequenced data underwent calibration, adapter trimming, identification of insert sequencing and UMI sequencing, alignment to the mouse-specific miRbase mature database and GRCm38 sequence, and read counting utilizing the “Primary Quantification” tool on GeneGlobe (https://geneglobe.qiagen.com/jp/). The miRDB database (https://www.mirdb.org) was utilized to predict the miRNAs targeting *Igf1r* gene and the pathway associated with IGF-1R expression.

### 2.8. Statistical analysis

All data are expressed as mean ± standard error (SEM) and were analyzed using Prism version 10.2.2 (MDF, Tokyo, Japan). Before the analysis for comparisons between sedentary and exercise groups, we checked the normality of raw data distribution using histograms. When the data were normally distributed, parametric tests (unpaired t-test) were used for statistical analyses. On the other hand, if we could not confirm that data were distributed normally, non-parametric tests (Mann–Whitney test) were used for statistical analyses. Statistical significance was set at *p* < 0.05. Log_2_ fold change in each miRNA and statistical significances were analyzed using the TCC-iDEGES-edgeR pipeline on R (https://www.R-project.org/, version 4.0.3) package TCC (version 1.30.0) [32] and edgeR (version 3.32.1).

## 3. Results

### 3.1. The effects of LC-HP diet on physiological and biochemical variables

In mice fed the LC-HP diet, blood glucose levels and the fat-to-body weight ratio were significantly lower than those fed the control diet (Table 2; *p* = 0.0027, *p* = 0.0022, respectively). Conversely, the kidney-to-body weight ratio in mice fed the LC-HP diet was significantly higher than in those fed the control diet (Table 2; *p* = 0.0022). However, there were no observed differences in the change ratio of body weight, the amounts of food intake, or β-ketone levels between mice fed the LC-HP diet and those fed the control diet (Table 2; *p* = 0.6366, *p* = 0.3019, *p* = 0.6980, respectively).

**Table 2.**
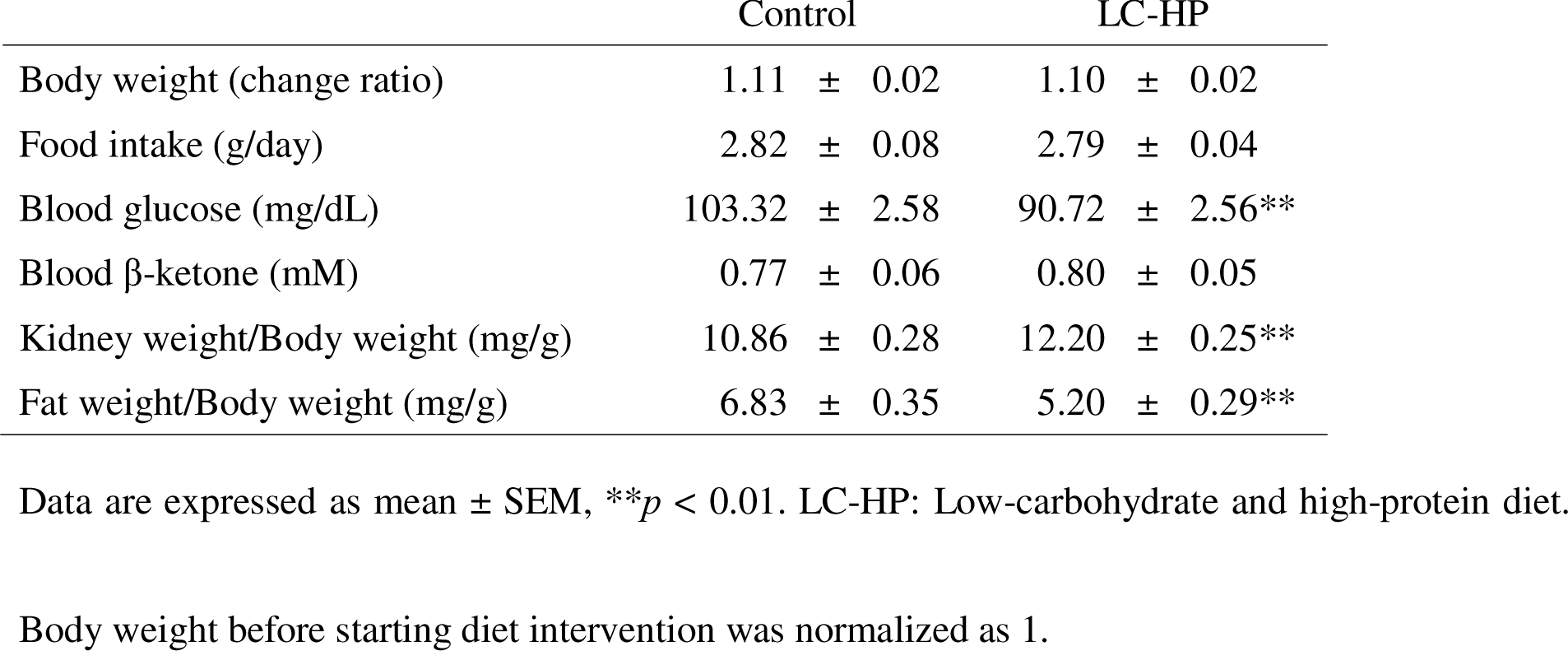
The effects of LC-HP diet on physiological and biochemical variables.

|  | Control | LC-HP |
| --- | --- | --- |
| Body weight (change ratio) | 1.11 ± 0.02 | 1.10 ± 0.02 |
| Food intake (g/day) | 2.82 ± 0.08 | 2.79 ± 0.04 |
| Blood glucose (mg/dL) | 103.32 ± 2.58 | 90.72 ± 2.56** |
| Blood β-ketone (mM) | 0.77 ± 0.06 | 0.80 ± 0.05 |
| Kidney weight/Body weight (mg/g) | 10.86 ± 0.28 | 12.20 ± 0.25** |
| Fat weight/Body weight (mg/g) | 6.83 ± 0.35 | 5.20 ± 0.29** |

Data are expressed as mean ± SEM, \*\* $p < 0.01$ . LC-HP: Low-carbohydrate and high-protein diet.
Body weight before starting diet intervention was normalized as 1.

### 3.2. Changes in working memory and hippocampal mRNA levels with feeding LC-HP diet

Although there was no difference in the number of arm entries (Fig. 1A; *p* = 0.5715), mice fed the LC-HP diet exhibited a lower correct ratio in the Y-maze test compared to those fed the control diet (Fig. 1B; *p* = 0.0037). Furthermore, the mRNA levels of *Igf1r* and *Dcx* were significantly downregulated in response to the LC-HP diet (Fig. 2A and C; *p* = 0.0321, *p* = 0.0425, respectively). On the other hand, *Bdnf* mRNA levels in the hippocampus of mice remained unaltered upon feeding the LC-HP diet (Fig. 2B; *p* = 0.2170).

**Fig. 1.**
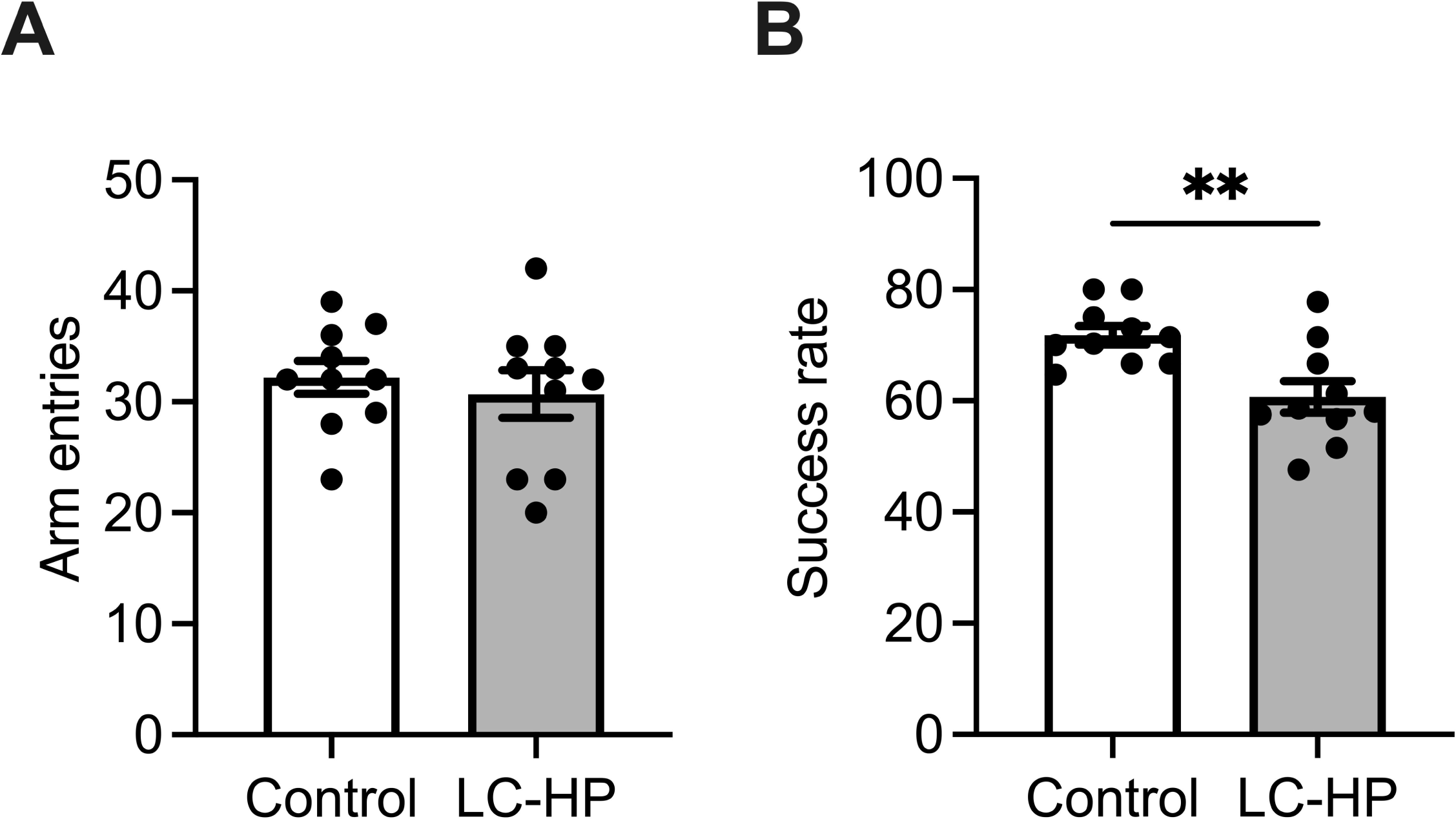
Effect of LC-HP diet on working memory function. The number of arm entries (A), and correct ratio (B) during Y-maze. White bars: mice fed control diet, and gray bars: mice fed LC-HP diet. Data are expressed as mean ± SEM, n = 10 mice for each group. \*\**p* < 0.01. LC-HP: low-carbohydrate and high-protein diet.

**Fig. 2.**
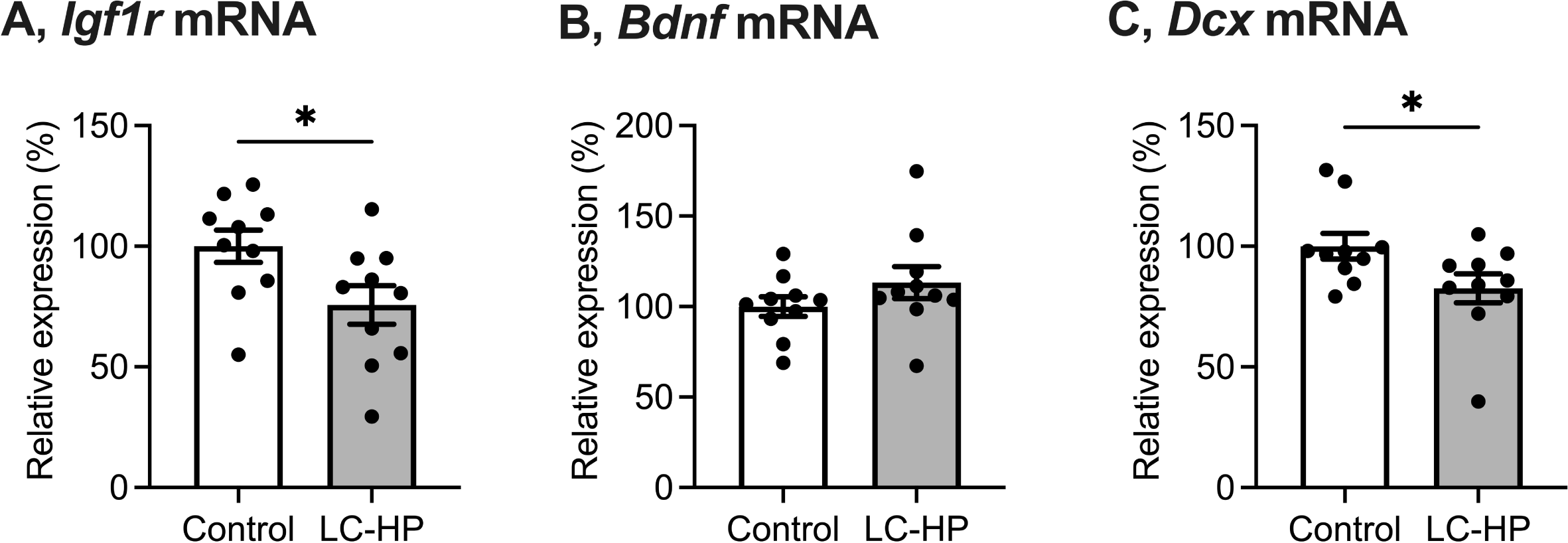
Effect of LC-HP diet on mRNA levels of *Igf1r* (A), *Bdnf* (B), and *Dcx* (C) in the hippocampus. White bars: mice fed control diet, and gray bars: mice fed LC-HP diet. Data are expressed as mean ± SEM, n = 10 mice for each group. \**p* < 0.05. LC-HP: low-carbohydrate and high-protein diet.

### 3.3. Quantification of miRNA in the hippocampus

There was no significant difference in the total reads of hippocampal miRNA between mice fed the LC-HP diet and those fed the control diet (Fig. 3A; *p* = 0.6129). Through miRNA sequencing, it was found that 44 kinds of hippocampal miRNA were significantly regulated by feeding LC-HP diet; among these, 17 were upregulated, and 27 were downregulated (Fig. 3B). The fold changes for these miRNAs are provided in Supplementary Tables 2 and 3. Notably, among the 17 miRNAs upregulated by the LC-HP diet, miR-497a-3p, miR-539-3p, miR-743a-3p, miR-3086-3p, and miR-7046-3p are predicted to target the *Igf1r* gene, as per miRDB (https://www.mirdb.org).

**Fig. 3.**
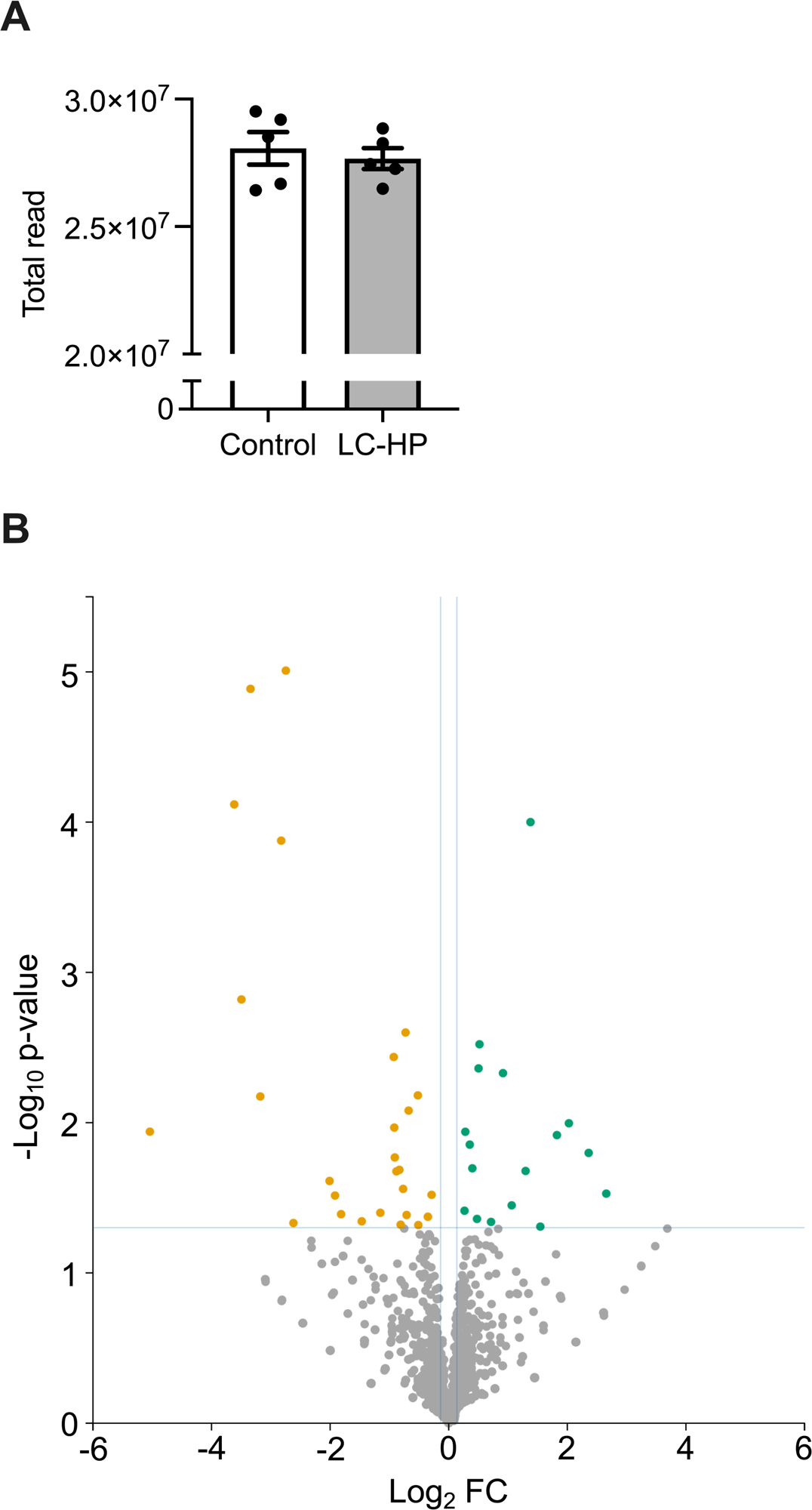
Total reads of miRNA by miR-seq in the hippocampus for each group (A). White bars: mice fed control diet, and gray bars: mice fed LC-HP diet. Data are expressed as mean ± SEM, n = 5 mice for each group. LC-HP: low-carbohydrate and high-protein diet. Significantly modulated miRNA in the hippocampus with feeding LC-HP diet (B).

### 3.4. Changes in miR-539-3p-related mRNA levels in the hippocampus with feeding LC-HP diets

The mice fed the LC-HP diet exhibited significantly lower hippocampal mRNA levels of low-density lipoprotein receptor-related protein 6 (*Lrp6*), a gene regulated by miR-539-3p, compared to those fed the control diet (Fig. 4A; *p* = 0.0261). In addition, hippocampal *Lrp6* mRNA levels showed a significant positive correlation with hippocampal *Igf1r* mRNA levels (Fig. 4E; *r* = 0.5990, *p* = 0.0053). Regarding other mRNAs regulated by miR-539-3p, hippocampal mRNA levels of a-kinase anchoring protein-3 (*Akap3*) were decreased with feeding the LC-HP diet, whereas no significant changes were observed in the mRNA levels of ring finger protein 2 (*Rnf2*) or a disintegrin and metalloproteinase with thrombospondin type 1 motif 5 (*Adamts5*) (Fig. 4B, C, and D; *p* = 0.0067, *p* = 0.7081, *p* = 0.9829, respectively).

**Fig. 4.**
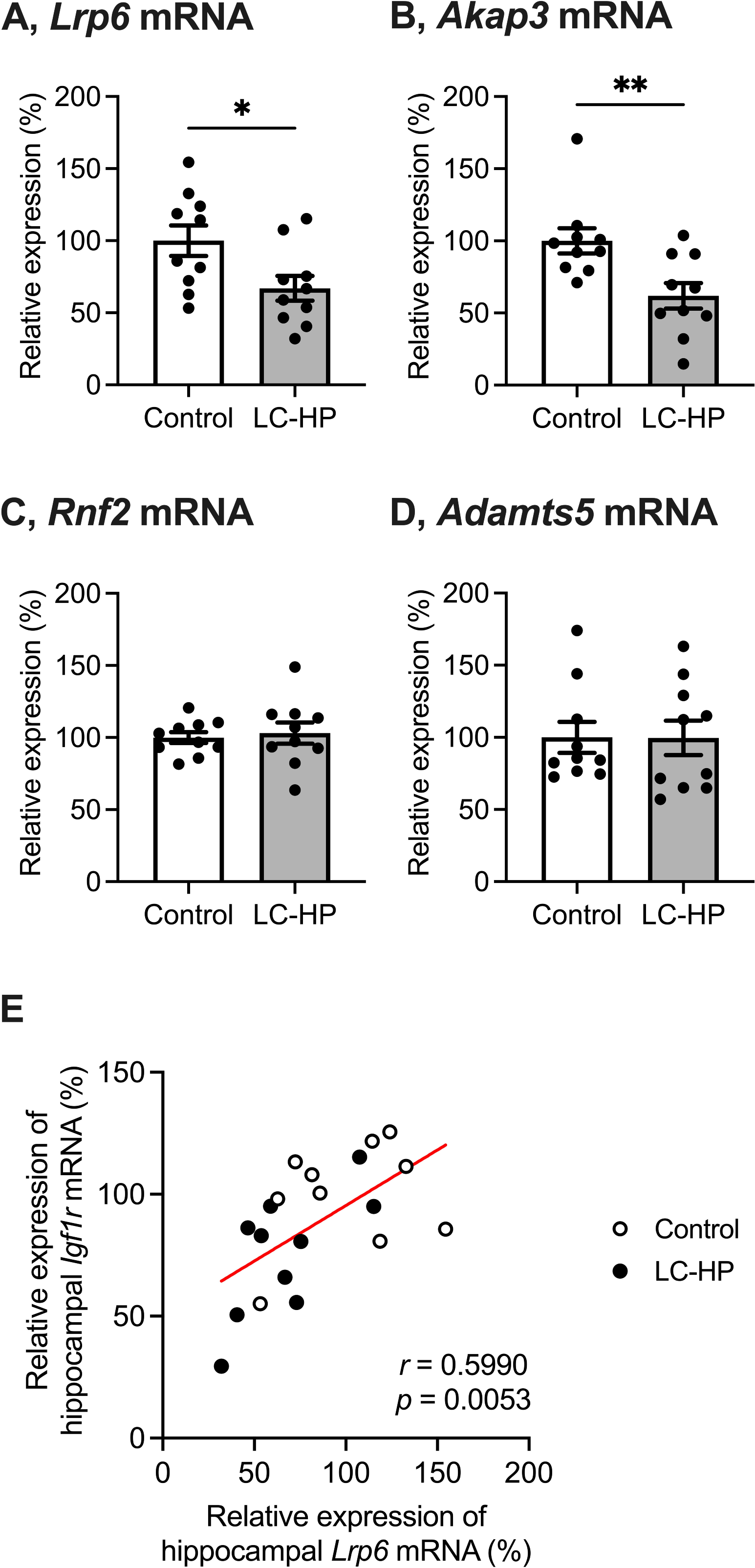
Effect of LC-HP diet on mRNA levels of *Lrp6* (A), *Akap3* (B), *Rnf2* (C), and *Adamts5* (D) in the hippocampus. White bars: mice fed control diet, and gray bars: mice fed LC-HP diet. Data are expressed as mean ± SEM, n = 10 mice for each group. \**p* < 0.05, \*\**p* < 0.01. LC-HP: low-carbohydrate and high-protein diet. The correlations between mRNA levels of *Lrp6* and *Igf1r* in the hippocampus (E). The line in the scatter diagram indicates significant correlation.

## Discussion

The current study investigated the effects of the LC-HP diet on hippocampal working memory and the levels of miRNA and mRNA in the hippocampus. Our findings reaffirm that feeding the LC-HP diet leads to a reduction in the correct ratio in the Y-maze, accompanied by decreased hippocampal *Igf1r* and *Dcx* mRNA levels. Furthermore, we observed significantly elevated levels of miR-539-3p and reduced levels of *Lrp6* mRNA in the hippocampus of mice fed the LC-HP diet compared to those fed the control diet.

Consistent with previous studies [2–4,7], the current study showed that mice fed the LC-HP diet exhibited significantly lower blood glucose levels, a reduced ratio of fat to body weight, and an increased ratio of kidney to body weight compared to those fed the control diet (Table 2). Additionally, four weeks of LC-HP diet feeding significantly suppressed working memory function in healthy mice (Fig. 1B), along with downregulation of hippocampal mRNA levels of *Igf1r* and *Dcx* (Fig. 2A and C). These results, consistent with those of a previous report [7], confirm the validity of the LC-HP diet utilized in our study. Given the crucial role of IGF-1R in regulating neurogenesis [12], it is assumed that IGF-1R-related neurogenesis in the hippocampus is involved in LC-HP diet-induced suppression of working memory based on our findings and prior reports [7]. Nevertheless, our study did not investigate morphological changes in the hippocampus associated with LC-HP diet feeding, indicating the necessity for further mechanism-based investigations.

Based on the miR-Seq results in our study, we identified five miRNAs (miR-497a-3p, miR-539-3p, miR-743a-3p, miR-3086-3p, and miR-7046-3p) predicted to target *Igf1r* gene according to miRDB (https://www.mirdb.org) among the upregulated hippocampal miRNAs associated with LC-HP diet feeding. Of particular interest was miR-539-3p, as it is also predicted to target pathways related to IGF-1R. Previous research has demonstrated that miR-539-3p downregulates *Lrp6* mRNA levels [33], and LRP6, in turn, regulates expressions and activation of IGF-1R [34,35]. LRP6 plays a critical role in Wnt/β-catenin signal transduction [36–38]. Further, LRP6/Wnt/β-catenin/IGF-1R signaling would contribute to cell growth [35]. Importantly, our study revealed a significant positive correlation between *Lrp6* and *Igf1r* mRNA in the hippocampus (Fig. 4E). Therefore, there is a possibility that the LC-HP diet suppresses hippocampal function mediated by the miR-539-3p/*Lrp6*/*Igf1r* axis in healthy mice.

MiR-539-3p targets not only *Igf1r* and *Lrp6* genes but also *Akap3*, *Rnf2*, and *Adamts5* genes, as reported in previous studies [33,39,40]. Our current study revealed that feeding the LC-HP diet downregulated hippocampal mRNA levels of *Akap3*, while no significant changes were observed in *Rnf2* or *Adamts5* mRNAs (Fig. 4B-D). AKAPs bind protein kinase A (PKA) and enhance neuroplasticity [41–43]. Therefore, the suppression of hippocampal function in healthy mice fed the LC-HP diet could be mediated by both the miR-539-3p/*Lrp6*/*Igf1r* axis and the miR-539-3p/*Akap*3 axis. However, our study did not investigate this aspect, indicating the need for further research in the future. Furthermore, miR-539-3p is also predicted to target *Bdnf* gene according to miRDB (https://www.mirdb.org). However, consistent with a previous report [7] and our current finding (Fig. 2B), hippocampal *Bdnf* mRNA levels remained unaltered when feeding the LC-HP diet. These findings suggest that miR-539-3p primarily targets *Igf1r*, *Lrp6*, and *Akap3* genes rather than *Rnf2*, *Adamts5*, and *Bdnf* genes in the mice fed the LC-HP diet.

Several miRNAs showing significant alterations in miR-Seq in our current study (Fig. 3B, supplementary Tables 2 and 3) have been implicated in neuroplasticity. For instance, upregulation of hippocampal miR144-3p has been observed in rodent models of schizophrenia [44], which is characterized by working memory deficit as a complication [45]. Additionally, a deficiency of miR-298-5p is associated with neurotoxicity [46]. Our results indicated increased levels of miR144-3p and decreased levels of miR-298-5p in the hippocampus of mice fed LC-HP diet (Fig. 3B and Supplementary Table 3), suggesting the involvement of these miRNAs in LC-HP diet-induced suppression of hippocampal function. Conversely, mice fed the LC-HP diet exhibited several changes in hippocampal miRNAs that could improve hippocampal function. Previous studies have reported that downregulation of miR-200c-3p and miR-335-3p contribute to improving hippocampal apoptosis and cognitive dysfunction [47,48]. However, in our study, downregulation of these miRNAs and suppression of working memory were concurrently observed in mice fed the LC-HP diet (Fig. 3B and Supplementary Table 3). Therefore, further mechanism-based investigations are warranted to elucidate the crucial pathways involved in LC-HP diet-induced suppression of hippocampal function.

The current study has some limitations. Firstly, we utilized only one specific composition of the LC-HP diet. Given that the sources of nutritional ingredients can influence the cognitive effects of dietary interventions [49,50], further validation is necessary. Secondly, our study focused solely on regulating hippocampal *Igf1r* mRNA, which does not conclusively establish the importance of hippocampal IGF-1R in LC-HP diet-induced suppression of working memory. Therefore, future investigations are warranted to elucidate this aspect further. Thirdly, we did not assess hippocampal protein levels or morphological changes in our study, which could provide valuable insights. Finally, our evaluation was confined to adaptations within the hippocampus. It is known that peripheral tissue-derived miRNAs packaged in exosomes can regulate hippocampal plasticity and function [51,52]. Therefore, peripheral organ-derived miRNAs may influence alterations in hippocampal miRNA profiles observed in our study. Exploring the crosstalk between peripheral organs and the hippocampus necessitates further investigation.

In conclusion, our study demonstrates that four weeks of LC-HP diet feeding leads to a decline in working memory function, accompanied by upregulation of hippocampal miR-539-3p and downregulation of hippocampal *Igf1r* and *Lrp6* mRNA levels in healthy mice. These findings suggest that the LC-HP diet suppresses hippocampal function in healthy mice through the miR-539-3p/*Lrp6*/*Igf1r* axis, indicating potential new targets for innovating and developing nutritional strategies to maintain and enhance hippocampal function.

## Authorship

**Takeru Shima**: Conceptualization, Methodology, Resources, Investigation, Writing–original draft, Writing–review & editing, Funding acquisition; **Hayate Onishi**: Investigation, Writing–review & editing; **Chiho Terashima**: Investigation, Writing–review & editing.

## Acknowledgements

We thank Mr. Yohei Morishita, Mr. Junpei Iijima, and Mr. Hirotaka Sutoh for their helpful technical assistance.

## Data availability

The datasets in the current study are available from the corresponding author on reasonable request.

## References

[1] Kinzig KP, Hargrave SL, Hyun J, Moran TH. Energy balance and hypothalamic effects of a high-protein/low-carbohydrate diet. Physiol Behav 2007;92:454–60. 10.1016/j.physbeh.2007.04.019.

[2] Higashida K, Terada S, Li X, Inoue S, Iida N, Kitai S, et al. Low-carbohydrate high-protein diet diminishes the insulin response to glucose load via suppression of SGLT-1 in mice. Biosci Biotechnol Biochem 2019;83:365–71. 10.1080/09168451.2018.1533803.

[3] Kostogrys RB, Franczyk-Zarów M, Maślak E, Topolska K. Effect of low carbohydrate high protein (LCHP) diet on lipid metabolism, liver and kidney function in rats. Environ Toxicol Pharmacol 2015;39:713–9. 10.1016/j.etap.2015.01.008.

[4] Kostogrys RB, Franczyk-Żarów M, Maślak E, Gajda M, Mateuszuk Ł, Jackson CL, et al. Low carbohydrate, high protein diet promotes atherosclerosis in apolipoprotein E/low-density lipoprotein receptor double knockout mice (apoE/LDLR−/−). Atherosclerosis 2012;223:327–31. 10.1016/j.atherosclerosis.2012.05.024.

[5] Foo SY, Heller ER, Wykrzykowska J, Sullivan CJ, Manning-Tobin JJ, Moore KJ, et al. Vascular effects of a low-carbohydrate high-protein diet. Proceedings of the National Academy of Sciences 2009;106:15418–23. 10.1073/pnas.0907995106.

[6] Galletly C, Moran L, Noakes M, Clifton P, Tomlinson L, Norman R. Psychological benefits of a high-protein, low-carbohydrate diet in obese women with polycystic ovary syndrome—A pilot study. Appetite 2007;49:590–3. 10.1016/j.appet.2007.03.222.

[7] Shima T, Yoshikawa T, Onishi H. Low-Carbohydrate and High-Protein Diet Suppresses Working Memory Function in Healthy Mice. J Nutr Sci Vitaminol (Tokyo) 2022;68:527–32. 10.3177/jnsv.68.527.

[8] Zhang J, Moats□Staats BM, Ye P, D’Ercole AJ. Expression of insulin□like growth factor system genes during the early postnatal neurogenesis in the mouse hippocampus. J Neurosci Res 2007;85:1618–27. 10.1002/jnr.21289.

[9] Chen BH, Ahn JH, Park JH, Song M, Kim H, Lee T-K, et al. Rufinamide, an antiepileptic drug, improves cognition and increases neurogenesis in the aged gerbil hippocampal dentate gyrus via increasing expressions of IGF-1, IGF-1R and p-CREB. Chem Biol Interact 2018;286:71–7. 10.1016/j.cbi.2018.03.007.

[10] Okamoto M, Yamamura Y, Liu Y, Min-Chul L, Matsui T, Shima T, et al. Hormetic effects by exercise on hippocampal neurogenesis with glucocorticoid signaling. Brain Plasticity 2015;1:149–58. 10.3233/BPL-150012.

[11] Yook JS, Rakwal R, Shibato J, Takahashi K, Koizumi H, Shima T, et al. Leptin in hippocampus mediates benefits of mild exercise by an antioxidant on neurogenesis and memory. Proceedings of the National Academy of Sciences 2019;116:10988–93. 10.1073/pnas.1815197116.

[12] Liu W, Ye P, O’Kusky JR, D’Ercole AJ. Type 1 insulin-like growth factor receptor signaling is essential for the development of the hippocampal formation and dentate gyrus. J Neurosci Res 2009;87:2821–32. 10.1002/jnr.22129.

[13] Moon HY, Becke A, Berron D, Becker B, Sah N, Benoni G, et al. Running-Induced Systemic Cathepsin B Secretion Is Associated with Memory Function. Cell Metab 2016;24:332–40. 10.1016/j.cmet.2016.05.025.

[14] Okamoto M, Inoue K, Iwamura H, Terashima K, Soya H, Asashima M, et al. Reduction in paracrine Wnt3 factors during aging causes impaired adult neurogenesis. FASEB J 2011;25:3570–82. 10.1096/fj.11-184697.

[15] Rossi C, Angelucci A, Costantin L, Braschi C, Mazzantini M, Babbini F, et al. Brain-derived neurotrophic factor (BDNF) is required for the enhancement of hippocampal neurogenesis following environmental enrichment. European Journal of Neuroscience 2006;24:1850–6. 10.1111/j.1460-9568.2006.05059.x.

[16] Lee J, Duan W, Mattson MP. Evidence that brain-derived neurotrophic factor is required for basal neurogenesis and mediates, in part, the enhancement of neurogenesis by dietary restriction in the hippocampus of adult mice. J Neurochem 2002;82:1367–75. 10.1046/j.1471-4159.2002.01085.x.

[17] Agrawal N, Dasaradhi PVN, Mohmmed A, Malhotra P, Bhatnagar RK, Mukherjee SK. RNA Interference: Biology, Mechanism, and Applications. Microbiology and Molecular Biology Reviews 2003;67:657–85. 10.1128/mmbr.67.4.657-685.2003.

[18] Bartel DP. MicroRNAs: Genomics, Biogenesis, Mechanism, and Function. Cell 2004;116:281–97. 10.1016/S0092-8674(04)00045-5.

[19] Ma Q, Zhang L, Pearce WJ. MicroRNAs in brain development and cerebrovascular pathophysiology. American Journal of Physiology-Cell Physiology 2019;317:C3–19. 10.1152/ajpcell.00022.2019.

[20] Shima T, Kawabata-Iwakawa R, Onishi H, Jesmin S, Yoshikawa T. Light-intensity exercise improves memory dysfunction with the restoration of hippocampal MCT2 and miRNAs in type 2 diabetic mice. Metab Brain Dis 2023;38:245–54. 10.1007/s11011-022-01117-y.

[21] Shima T, Kawabata-Iwakawa R, Onishi H, Jesmin S, Yoshikawa T. Four weeks of light-intensity exercise enhances empathic behavior in mice: The possible involvement of BDNF. Brain Res 2022;1787:147920. 10.1016/j.brainres.2022.147920.

[22] Cannataro R, Cione E. Diet and miRNA: Epigenetic Regulator or a New Class of Supplements? MicroRNA 2022;11:89–90. 10.2174/2211536611666220510111711.

[23] Que E, James KL, Coffey AR, Smallwood TL, Albright J, Huda MN, et al. Genetic architecture modulates diet-induced hepatic mRNA and miRNA expression profiles in Diversity Outbred mice. Genetics 2021;218. 10.1093/genetics/iyab068.

[24] Yu Y, Zhang J, Wang J, Sun B. MicroRNAs: The novel mediators for nutrient-modulating biological functions. Trends Food Sci Technol 2021;114:167–75. 10.1016/j.tifs.2021.05.028.

[25] Li X, Lian F, Liu C, Hu K-Q, Wang X-D. Isocaloric Pair-Fed High-Carbohydrate Diet Induced More Hepatic Steatosis and Inflammation than High-Fat Diet Mediated by miR-34a/SIRT1 Axis in Mice. Sci Rep 2015;5:16774. 10.1038/srep16774.

[26] Su Y, Jiang X, Li Y, Li F, Cheng Y, Peng Y, et al. Maternal Low Protein Isocaloric Diet Suppresses Pancreatic β-Cell Proliferation in Mouse Offspring via miR-15b. Endocrinology 2016;157:4782–93. 10.1210/en.2016-1167.

[27] Ramzan F, D’Souza RF, Durainayagam BR, Milan AM, Roy NC, Kruger MC, et al. Inflexibility of the plasma miRNA response following a high-carbohydrate meal in overweight insulin-resistant women. Genes Nutr 2020;15:2. 10.1186/s12263-020-0660-8.

[28] Zhang J, Zhang F, Didelot X, Bruce KD, Cagampang FR, Vatish M, et al. Maternal high fat diet during pregnancy and lactation alters hepatic expression of insulin like growth factor-2 and key microRNAs in the adult offspring. BMC Genomics 2009;10:478. 10.1186/1471-2164-10-478.

[29] Xu C, Niu J-J, Zhou J-F, Wei Y-S. MicroRNA-96 is responsible for sevoflurane-induced cognitive dysfunction in neonatal rats via inhibiting IGF1R. Brain Res Bull 2019;144:140–8. 10.1016/j.brainresbull.2018.09.001.

[30] Shi Z, Wang X, Qian X, Tao T, Wang L, Chen Q, et al. MiRNA-181b suppresses IGF-1R and functions as a tumor suppressor gene in gliomas. RNA 2013;19:552–60. 10.1261/rna.035972.112.

[31] Feng S-J, Zhang X-Q, Li J-T, Dai X-M, Zhao F. miRNA-223 regulates ischemic neuronal injury by targeting the type 1 insulin-like growth factor receptor (IGF1R). Folia Neuropathol 2018;56:49–57. 10.5114/fn.2018.74659.

[32] Sun J, Nishiyama T, Shimizu K, Kadota K. TCC: An R package for comparing tag count data with robust normalization strategies. BMC Bioinformatics 2013;14. 10.1186/1471-2105-14-219.

[33] Tripathi A, John AA, Kumar D, Kaushal SK, Singh DP, Husain N, et al. MiR-539-3p impairs osteogenesis by suppressing Wnt interaction with LRP-6 co-receptor and subsequent inhibition of Akap-3 signaling pathway. Front Endocrinol (Lausanne) 2022;13:1–19. 10.3389/fendo.2022.977347.

[34] Singh R, De Aguiar RB, Naik S, Mani S, Ostadsharif K, Wencker D, et al. LRP6 Enhances Glucose Metabolism by Promoting TCF7L2-Dependent Insulin Receptor Expression and IGF Receptor Stabilization in Humans. Cell Metab 2013;17:197–209. 10.1016/j.cmet.2013.01.009.

[35] Aggelidakis J, Berdiaki A, Nikitovic D, Papoutsidakis A, Papachristou DJ, Tsatsakis AM, et al. Biglycan regulates MG63 osteosarcoma cell growth through a Lpr6/β-catenin/IGFR-IR signaling axis. Front Oncol 2018;8:1–15. 10.3389/fonc.2018.00470.

[36] Singh HD, Ma J xing, Takahashi Y. Distinct roles of LRP5 and LRP6 in Wnt signaling regulation in the retina. Biochem Biophys Res Commun 2021;545:8–13. 10.1016/j.bbrc.2021.01.068.

[37] Ren Q, Chen J, Liu Y. LRP5 and LRP6 in Wnt Signaling: Similarity and Divergence. Front Cell Dev Biol 2021;9:1–11. 10.3389/fcell.2021.670960.

[38] Jeong W, Jho E. Regulation of the Low-Density Lipoprotein Receptor-Related Protein LRP6 and Its Association With Disease: Wnt/β-Catenin Signaling and Beyond. Front Cell Dev Biol 2021;9:1–14. 10.3389/fcell.2021.714330.

[39] Wu X-J, Xie Y, Gu X-X, Zhu H-Y, Huang L-X. LncRNA XIST promotes mitochondrial dysfunction of hepatocytes to aggravate hepatic fibrogenesis via miR-539-3p/ADAMTS5 axis. Mol Cell Biochem 2023;478:291–303. 10.1007/s11010-022-04506-0.

[40] Yue F, Peng K, Zhang L, Zhang J. Circ_0004104 Accelerates the Progression of Gastric Cancer by Regulating the miR-539-3p/RNF2 Axis. Dig Dis Sci 2021;66:4290–301. 10.1007/s10620-020-06802-5.

[41] Dell’Acqua ML, Smith KE, Gorski JA, Horne EA, Gibson ES, Gomez LL. Regulation of neuronal PKA signaling through AKAP targeting dynamics. Eur J Cell Biol 2006;85:627–33. 10.1016/j.ejcb.2006.01.010.

[42] Bhattacharyya S, Biou V, Xu W, Schlüter O, Malenka RC. A critical role for PSD-95/AKAP interactions in endocytosis of synaptic AMPA receptors. Nat Neurosci 2009;12:172–81. 10.1038/nn.2249.

[43] Gorshkov K, Mehta S, Ramamurthy S, Ronnett G V, Zhou F, Zhang J. AKAP-mediated feedback control of cAMP gradients in developing hippocampal neurons. Nat Chem Biol 2017;13:425–31. 10.1038/nchembio.2298.

[44] Pan B, Wang Y, Shi Y, Yang Q, Han B, Zhu X, et al. Altered expression levels of miR-144-3p and ATP1B2 are associated with schizophrenia. The World Journal of Biological Psychiatry 2022;23:666–76. 10.1080/15622975.2021.2022757.

[45] Van Snellenberg JX, Girgis RR, Horga G, van de Giessen E, Slifstein M, Ojeil N, et al. Mechanisms of Working Memory Impairment in Schizophrenia. Biol Psychiatry 2016;80:617–26. 10.1016/j.biopsych.2016.02.017.

[46] Wei X, Xu S, Chen L. LncRNA Neat1/miR-298-5p/Srpk1 Contributes to Sevoflurane-Induced Neurotoxicity. Neurochem Res 2021;46:3356–64. 10.1007/s11064-021-03436-5.

[47] Raihan O, Brishti A, Molla MR, Li W, Zhang Q, Xu P, et al. The Age-dependent Elevation of miR-335-3p Leads to Reduced Cholesterol and Impaired Memory in Brain. Neuroscience 2018;390:160–73. 10.1016/j.neuroscience.2018.08.003.

[48] Du Y, Chi X, An W. Downregulation of microRNA-200c-3p reduces damage of hippocampal neurons in epileptic rats by upregulating expression of RECK and inactivating the AKT signaling pathway. Chem Biol Interact 2019;307:223–33. 10.1016/j.cbi.2019.04.027.

[49] Muth AK, Park SQ. The impact of dietary macronutrient intake on cognitive function and the brain. Clinical Nutrition 2021;40:3999–4010. 10.1016/j.clnu.2021.04.043.

[50] Liu C, Meng Q, Zu C, Wei Y, Su X, Zhang Y, et al. Dietary low- and high-quality carbohydrate intake and cognitive decline: A prospective cohort study in older adults. Clinical Nutrition 2023;42:1322–9. 10.1016/j.clnu.2023.06.021.

[51] Wang J, Li L, Zhang Z, Zhang X, Zhu Y, Zhang C, et al. Extracellular vesicles mediate the communication of adipose tissue with brain and promote cognitive impairment associated with insulin resistance. Cell Metab 2022;34:1264–1279.e8. 10.1016/j.cmet.2022.08.004.

[52] Zhang Y, Xu C. Effects of exosomes on adult hippocampal neurogenesis and neuropsychiatric disorders. Mol Biol Rep 2022;49:6763–77. 10.1007/s11033-022-07313-4.

