## Supplementary Tables for "Feeding a low-carbohydrate and high-protein diet diminishes working memory in healthy mice: possible involvement of miR-539-3p/*Lrp6*/*Igf1r* axis in the hippocampus"

Supplementary Table 1. Primer sequences

| Gene | Primer | Sequence |
| --- | --- | --- |
| *Igf-1r* | F | CCAGGGCTTGTCCAACGAGCA |
|  | R | TCCCGGAAGCCAGGCTCCAT |
| *Bdnf* | F | GATGAGGACCAGAAGGTTCG |
|  | R | GATTGGGTAGTTCGGCATTG |
| *Trkb* | F | ATCCTGGTGGCCGTGAAG |
|  | R | TGAAAGTCCTTGCGAGCATTG |
| *Creb1* | F | TCAGCCGGGTACTACCATTC |
|  | R | TCTCTTGCTGCTTCCCTGTT |
| *Dcx* | F | GAGTGCGCTACATTTATACCATTG |
|  | R | TGACATTCTTGGTGTACTCAACCT |
| *Lrp6* | F | CATGGACATCCAAGTGCTGA |
|  | R | TTGTCCTCCTCGCATGGT |
| *Akap3* | F | AACAGAAAATTACTAAGCACCAACG |
|  | R | TTGGTGTCTTCACTATCCCTAAGTC |
| *Rnf2* | F | ATGCCCTACCTGTCGGAAAA |
|  | R | TCTTCAGCCCCTCCTCAATG |
| *Adamts5* | F | GCATCCAAGCCCTGGTCCAAAT |
|  | R | GGTGGCATCGTAGGTCTGTCCT |
| *β-actin* | F | TATGCCAACACAGTGCTGTCTGG |
|  | R | TACTCCTGCTTGCTGATCCACAT |

Supplementary Table 2. Fold change of significant upregulated miRNA in the hippocampus with Feeding LC-HP diet

| Name | Fold change | P-value |
| --- | --- | --- |
| mmu-miR-467e-3p | 6.306 | 0.02966 |
| mmu-miR-7000-3p | 5.133 | 0.01590 |
| mmu-miR-7046-3p | 4.074 | 0.01009 |
| mmu-miR-3086-3p | 3.541 | 0.01210 |
| mmu-miR-743a-3p | 2.918 | 0.04918 |
| mmu-miR-539-3p | 2.600 | 0.00010 |
| mmu-miR-144-3p | 2.456 | 0.02097 |
| mmu-miR-6970-3p | 2.088 | 0.03552 |
| mmu-miR-144-5p | 1.886 | 0.00468 |
| mmu-miR-7021-5p | 1.640 | 0.04575 |
| mmu-miR-497a-3p | 1.433 | 0.00301 |
| mmu-miR-19b-3p | 1.417 | 0.00436 |
| mmu-miR-3101-5p | 1.392 | 0.04371 |
| mmu-miR-3069-3p | 1.318 | 0.02013 |
| mmu-miR-101a-3p | 1.279 | 0.01400 |
| mmu-miR-551b-3p | 1.214 | 0.01150 |
| mmu-miR-218-2-3p | 1.202 | 0.03857 |

Supplementary Table 3. Fold change of significant downregulated miRNA in the hippocampus with Feeding LC-HP diet

| Name | Fold change | P-value |
| --- | --- | --- |
| mmu-miR-216b-3p | -32.910 | 0.01148 |
| mmu-miR-216a-3p | -12.270 | 0.00008 |
| mmu-miR-217-3p | -11.279 | 0.00151 |
| mmu-miR-217-5p | -10.142 | 0.00001 |
| mmu-miR-489-3p | -9.054 | 0.00670 |
| mmu-miR-216a-5p | -7.087 | 0.00013 |
| mmu-miR-3547-3p | -6.716 | 0.00001 |
| mmu-miR-344e-5p | -6.144 | 0.04652 |
| mmu-miR-1247-5p | -4.027 | 0.02444 |
| mmu-miR-365-1-5p | -3.777 | 0.03055 |
| mmu-miR-7662-3p | -3.519 | 0.04065 |
| mmu-miR-7032-3p | -2.764 | 0.04533 |
| mmu-miR-486a-3p | -2.225 | 0.03978 |
| mmu-miR-205-5p | -1.902 | 0.00366 |
| mmu-miR-135a-1-3p | -1.890 | 0.01078 |
| mmu-miR-539-5p | -1.878 | 0.01704 |
| mmu-miR-298-5p | -1.842 | 0.02110 |
| mmu-miR-1955-3p | -1.781 | 0.02061 |
| mmu-miR-219b-5p | -1.754 | 0.04773 |
| mmu-miR-6540-3p | -1.705 | 0.02759 |
| mmu-miR-335-3p | -1.657 | 0.00251 |
| mmu-miR-200c-3p | -1.637 | 0.04115 |
| mmu-miR-490-5p | -1.598 | 0.00831 |
| mmu-miR-770-3p | -1.434 | 0.00658 |
| mmu-miR-223-3p | -1.427 | 0.04800 |
| mmu-miR-3572-3p | -1.276 | 0.04227 |
| mmu-miR-540-3p | -1.222 | 0.03022 |
